# Creation and Validation of a Proteome-Wide Yeast Library for Protein Detection and Analysis

**DOI:** 10.1101/2025.01.14.632936

**Authors:** Din Baruch, Ioannis Tsirkas, Ehud Sass, Benjamin Dubreuil, Yeynit Asraf, Amir Aharoni, Maya Schuldiner, Ofir Klein

## Abstract

A significant challenge in cell biology is to uncover the function of uncharacterized proteins. Surprisingly a quarter of the proteome is still poorly understood even in the most well studied model organisms. Systematic methodologies, including the use of tagged protein collections, have emerged as a powerful approach to address this gap. Despite the availability of proteome-wide collections featuring various fused proteins, the impact of tag size on protein function highlighted the need for using minimally disruptive tags for functional genomic studies. To rise to this challenge, we have created a proteome-wide collection of yeast strains in which proteins are N-terminally tagged with the Hemagglutinin (HA) epitope. The library leverages the compact size of the HA tag to minimize drawbacks associated with larger tags while enabling efficient functional analysis. We showcase the potential uses of our library for systematically evaluating protein size, abundance and localization using an *in vivo* labeling approach. Our characterization underscores the potential utility of a proteome-wide HA-tagged library in revealing novel aspects of cell biology, providing an additional powerful tool for functional genomics.

## Introduction

In the post-genomic era, a paramount challenge still lies in mapping the functions of all uncharacterized proteins (Rocha et al., 2023). Despite the fact that both the yeast and human genomes were sequenced decades ago (Goffeau et al., 1996; International Human Genome Sequencing Consortium, 2004), there are still thousands of proteins, constituting up to a quarter of the proteome, with unknown or poorly understood functions (Cohen et al., 2022; Eisenhaber, 2012; Peña-Castillo and Hughes, 2007; Rocha et al., 2023; Wood et al., 2019). Uncovering the function of uncharacterized proteins can be approached either one protein at a time or through systematic, high-throughput, strategies, also termed functional genomics. Among the various systematic methodologies, pull-down assays for interactor mapping, and localization studies are commonly employed. These methods often rely on the labeling of the proteins by genetically encoded tags.

An established tool allowing for the simultaneous study of thousands of proteins is the use of libraries – collections of tagged proteins, where each protein is genetically fused to a tag of interest, one protein at a time. While systematic libraries have been created for several organisms (just a few as an example: (Bischof et al., 2013; Cho et al., 2022)) the most utilized eukaryotic model organism in library creation is the baker’s yeast *Saccharomyces cerevisiae* (yeast from now on). Yeast is a popular and well-studied model organism for functional genomics due to the availability of extensive genetic and genomic tools, and since approximately two-thirds of its proteome is conserved to humans (Cohen et al., 2022).

Although many yeast libraries have been created with different tags, including several fluorescent proteins (FPs) (Huh et al., 2003; Meurer et al., 2018; Weill et al., 2018), their large tag size can disrupt protein folding, complex assembly, stability, targeting and function, hindering their use for functional genomic assays.

Several small tags have been engineered, minimizing the impacts on the structure, function, and localization of the tagged protein. particularly useful one is the Hemagglutinin (HA) tag, just nine amino acids in length. The HA tag is not only effective, like other small tags, for immunoprecipitation (IP) studies, but was also recently used for *in vivo* visualization in combination with a newly developed genetically encoded single-chain fragment variable (scFv) fused to a FP (Murakawa et al., 2022; Tsirkas et al., 2022; Zhao et al., 2019). Recognizing these advantages of the HA tag, we created a proteome-wide yeast library where each protein is N-terminally (N’) tagged with HA. For library creation we utilized the SWAp-Tag (SWAT) strategy which revolutionized the speed and capacity to create “tailor made” yeast libraries (Meurer et al., 2018, 2018; Weill et al., 2018; Yofe et al., 2016). We demonstrate the HA collection efficacy in functionally diverse applications including the analysis of protein size, abundance and localization, highlighting the beneficial effect of the HA tag in comparison to the green fluorescent protein (GFP) tag on native protein localization. Overall, our efforts provide the yeast community with a new and powerful tool for functional genomics, that will be freely distributed.

## Results

### Creation and Validation of an N’-HA Tagged Library

Until now, yeast proteome-wide collections that were constructed for protein visualization (Huh et al., 2003; Meurer et al., 2018; Weill et al., 2018) or purification (Ghaemmaghami et al., 2003) relied on large tags such as GFP, mCherry, mScarlet-I, mNeonGreen and TAP (Tandem Affinity Purificaiton), respectively. These tags, while effective, may disrupt protein function due to their large size. The creation of our proteome-wide N’ HA-tagged yeast library was motivated by the need for minimally disruptive yet effective tags for functional genomics.

To create the library we used the SWAT strategy which employs an acceptor library with an interchangeable cassette that can be replaced with a tag of choice by homologous recombination to rapidly and efficiently generate new yeast libraries (Yofe et al., 2016) (Figure 1a). The resulting library comprises of a systematic collection of proteins N’ fused to a 3xHA epitope tag, positioned under the control of the *TEF1* constitutive promoter and a nourseothricin (Nat) resistance cassette, in *MATa*mating type, ensuring compatibility for downstream applications (Figure 1a).

**Figure 1.**
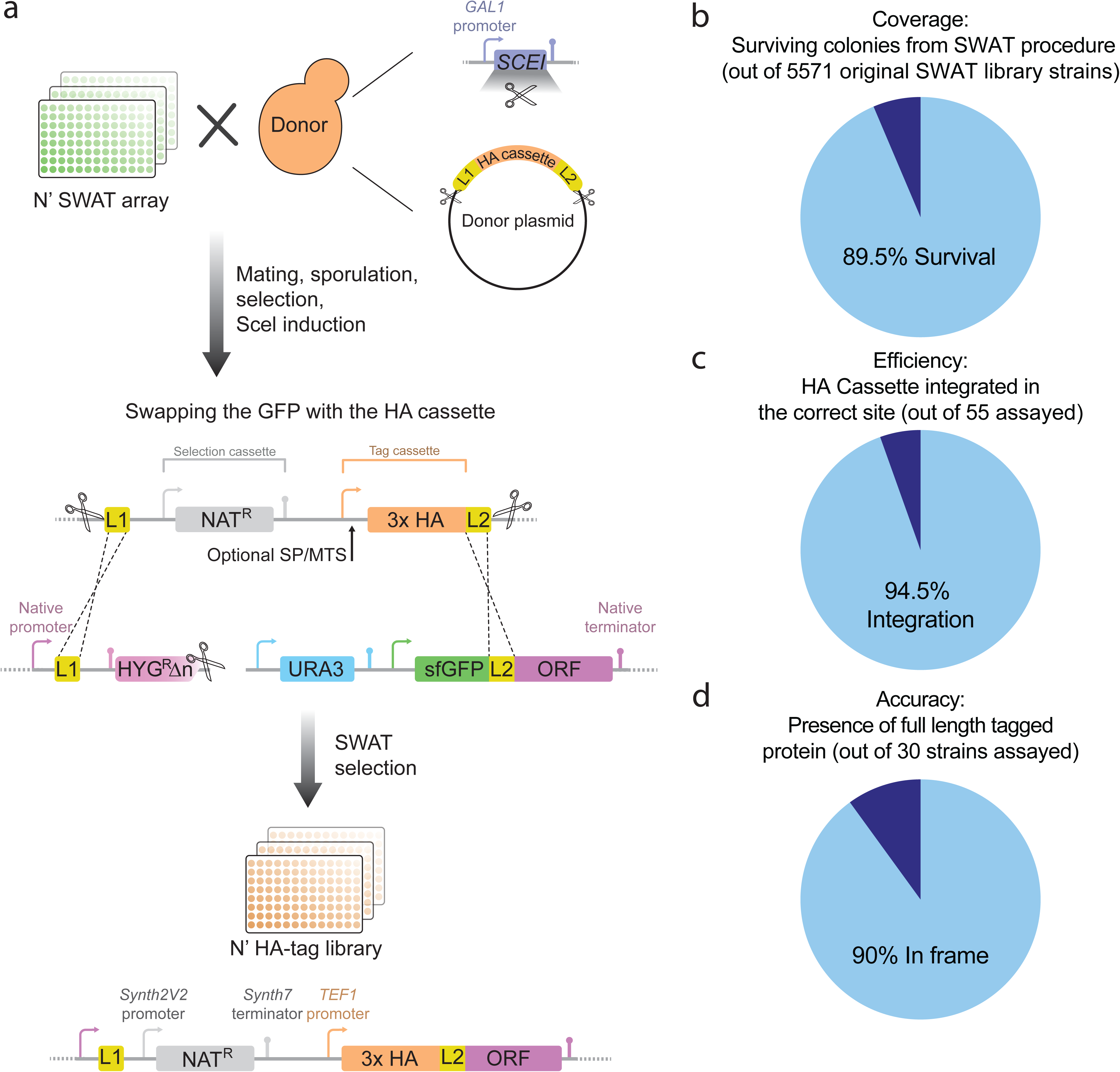
Creation and validation of an HA-tag yeast library. **(a)** An illustration of the HA library creation. The acceptor N’-SWAT library was crossed with a donor strain harboring a donor plasmid, encoding for the HA cassette and a galactose inducible I-SceI restriction enzyme. An automated mating and selection procedure created an intermediate library carrying all genetic traits. Induction of I-SceI expression resulted in double- strand breaks in the donor plasmid and the genomic SWAT module, leading to homologous recombination between the L1 and L2 linkers. ‘Swapped’ colonies were selected, and a library featuring the new tag was created. **(b)** Assessment of the efficiency of the SWAT procedure (coverage) was performed by measuring the number of colonies that passed all the automated mating, selection, and SWAT procedures. A pie chart represents that 89.5% of yeast strains from the original library survived. **(c)** Assessment of the HA cassette SWAT efficiency was performed by check PCR upstream to randomly chosen genes. The pie chart demonstrates that 94.5% of genes had a successful integration (n=55). **(d)** Assessment of the efficiency of in-frame integration (accuracy) was performed by evaluating the molecular weight (Mw) of tagged proteins. ‘The pie chart demonstrates that, for randomly chosen soluble proteins analyzed by SDS-PAGE, 90% of proteins run at the expected MW and therefore we assume that they represent in-frame integration of the HA tag (n=30).

In line with previous SWAT-based generated libraries (Weill et al., 2018) we had a very high efficiency of the process with 89.5% of yeast strains from the original library surviving the procedure (Figure 1b, supplementary table S1). Moreover, 94.5% of the 55 sampled strains exhibited successful integration of the HA tag, as confirmed by PCR analysis (Figure 1c). The accuracy of in-frame integration was further assessed by SDS-PAGE, revealing that 90% of the 30 randomly selected soluble proteins were detected at the expected molecular weight of the fusion protein (Figure 1d and supplementary figure 1). Furthermore, all 36 colonies tested for the loss of GFP signal from the original acceptor cassette, had indeed lost it. Altogether, we approximate that out of 5571 yeast strains present in the original SWAT library, a minimum of 4486 strains are *bona fide* HA-tagged proteins in our collection. This high coverage underscores the effectiveness of the SWAT procedure as a method to create novel libraries. More importantly, it makes our N’ HA-tagged library a powerful tool for systematic, proteome-wide, functional genomic studies. This library can provide a reliable resource for downstream systematic applications such as tracking protein size, abundance and localization, enabling comprehensive analysis across various experimental conditions.

### Rapid examination of post-translational modifications using the N’ HA-tagged strains

To demonstrate the usability of the new library for western blot analysis, we chose to examine the glycosylation state of ten known glycosylated proteins using PNGase digestion for glycan removal. This analysis is based on the addition of the PNGase enzyme to the lysate. This results in the enzymatic digestion of the glycan tree conjugated to glycosylated proteins, resulting in a reduction of their molecular weight. This can easily be visualized by a SDS-PAGE gel. The power of our collection is that instead of requiring an antibody for each protein one wishes to detect, it is possible to assay multiple proteins at once using a single, anti-HA, antibody. To showcase the ease of use we chose ten known glycosylated proteins of various molecular weights: Npp1, Sga1, Sag1, Pma2, Ecm30, Fmn1, Nce102, Vel1, Cwp1 and Ynl194c (Cao et al., 2014; Yeo et al., 2016; Zielinska et al., 2012) (Figure 2). Six out of the 10 showed a clear molecular weight change after PNGase treatment (Npp1, Sga1, Fmn1, Vel1, Cwp1, Ynl194c). Out of the 4 undetected proteins, 3 did not have a clear band: Pma2, Ecm30 and Sag1. Pma2 is a multi-pass membrane protein with 9-10 predicted transmembrane domains (Weill et al., 2019) and Ecm30 is a high-molecular weight protein of 150 kDa, and therefore are difficult to resolve under the same conditions as the other proteins. Sag1 is an alpha-agglutinin, and therefore while it is expressed in the HA library under a constitutive promoter, it is likely to be unstable in MATα cells under non-mating conditions. Lastly, Nce102 presented a similar band pattern with and without PNGase treatment at a molecular weight calculated for the unmodified protein. Hence, we might not have assayed this protein under conditions where it is in its glycosylated form.

**Figure 2.**
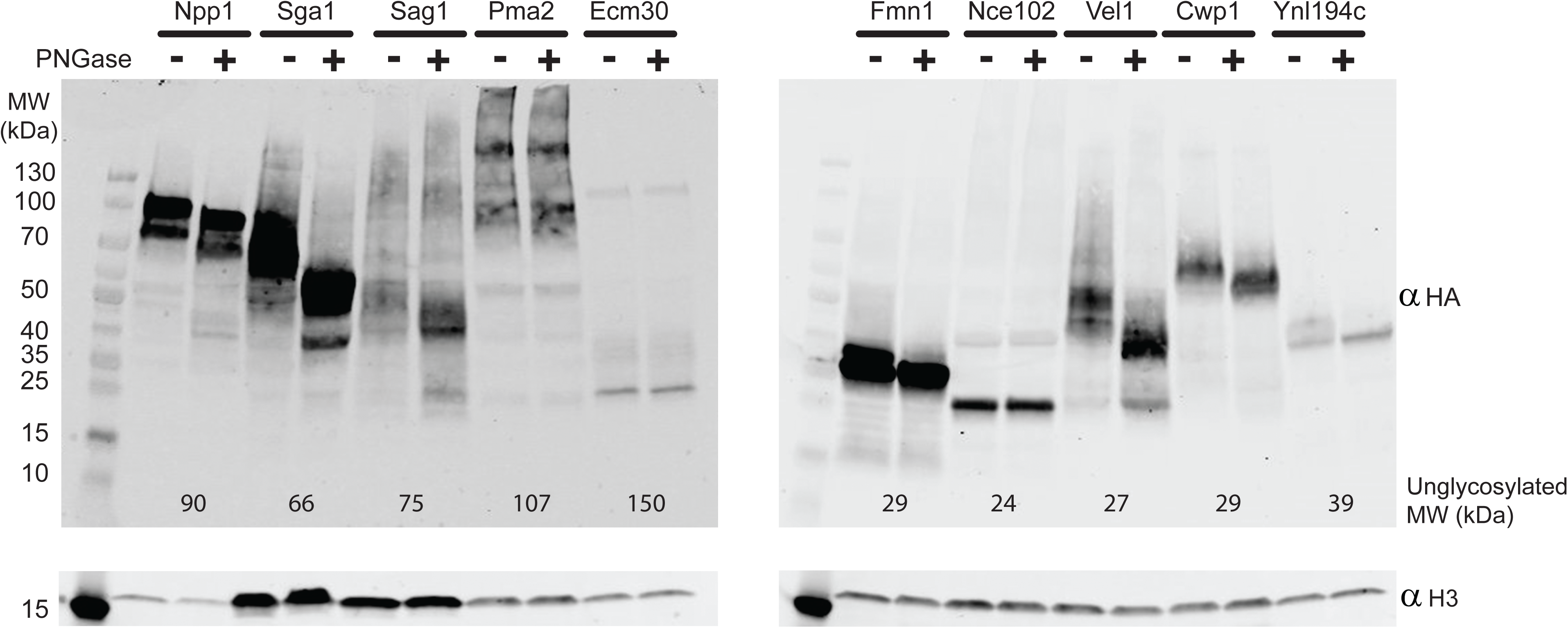
**Evaluation of the glycosylation state of HA-tagged proteins.** Whole cell lysates from colonies of 10 known glycoproteins were treated with the pan-glycan removing enzyme, PNGase (+), or DDW (-) as control. Their molecular weight (Mw) was subsequently resolved by western blot analysis using anti HA antibody. The calculated Mw of each protein, including the size of the 3xHA tag and L2 linker (4.9 KDa) is presented at the bottom of their respective lanes. Anti histone H3 was used as a loading control.

Regardless, our assay demonstrates how the HA-tagged library facilitates rapid and systematic analysis of post-translational modifications across a diverse set of proteins, enabling insights into glycosylation and other modifications. In a broader sense, these results highlight how the availability of a full proteome library of HA tagged proteins enables rapid size analysis for a selected set of proteins.

### Systematic Quantification of Protein Abundance Using N’-HA Strains

To demonstrate the versatility of the N’-HA library in systematic assays of protein abundance, we quantified protein levels independently of promoter activity. This quantification is essential for identifying variations in protein expression that are independent of promoter activity but rather influenced by factors such as RNA stability and protein degradation rates.

For this analysis we chose a set of proteins selected for their varying expression levels in the *Nop1*-GFP library (Weill et al., 2018) despite being regulated by the same, constitutive, promoter. In the N’-HA library, these proteins were also expressed from a constitutive *TEF1* promoter.

To measure their abundance we utilized high-throughput quantitative dot blotting (Figure 3a,b). Calibrating the signal to the total amount of extracted proteins our analysis revealed clear differences in the abundance of the selected proteins, indicating that the N’ HA-tagged library enables effective detection of variations in protein levels (Figure 3c). These results highlight the N’-HA library potential for reproducible and quantitative systematic assays for the analysis of cellular protein expression.

**Figure 3.**
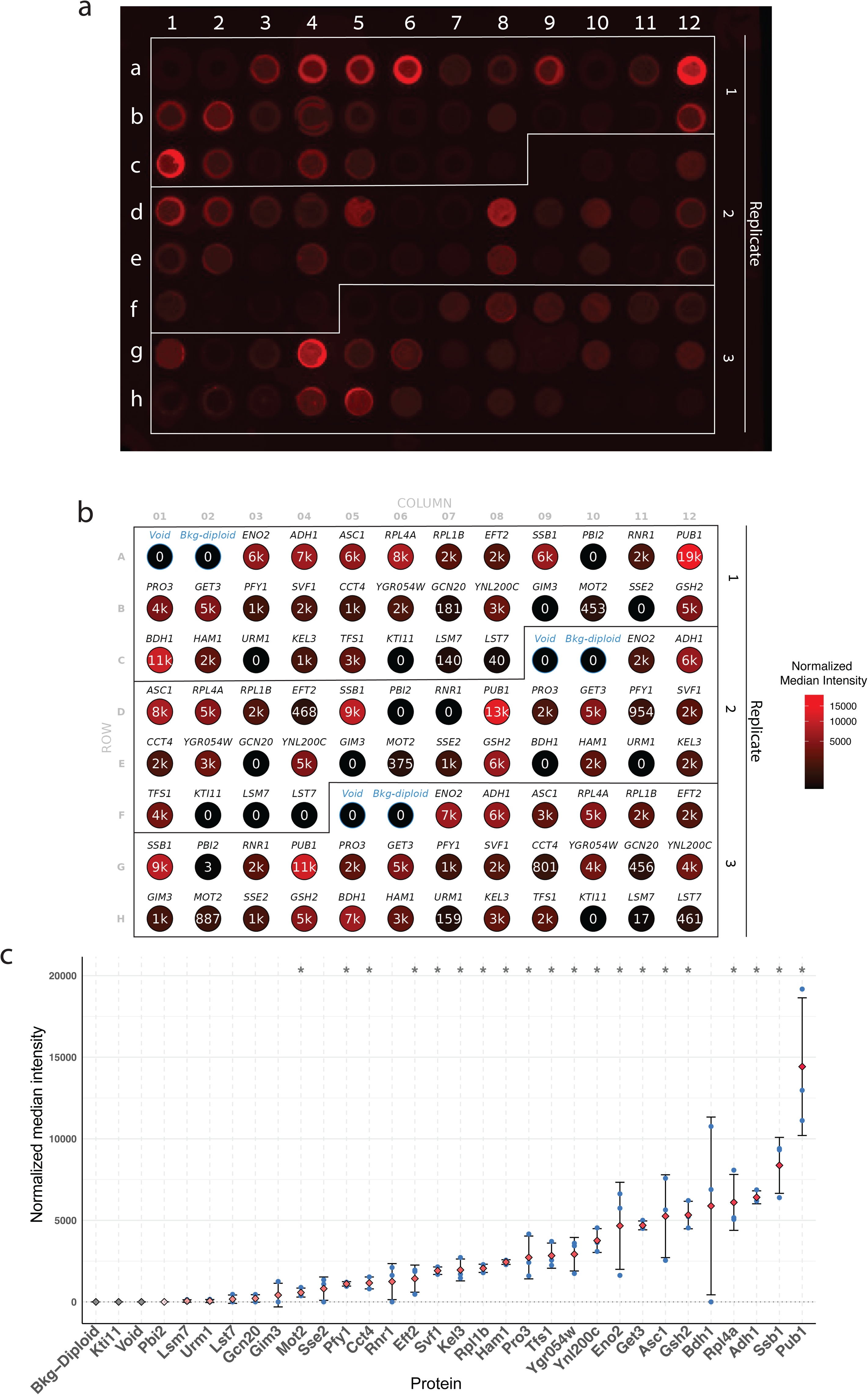
Systematic quantification of protein abundance using the N’ HA strains. (a) Image of a 96-well dot blot with an anti-HA antibody, showing three replicates for each of 30 yeast strains and controls, one without cells and one using a strain not expressing any HA tagged protein. **(b)** Well annotation with median intensity values normalized to BCA protein concentrations. Circle color intensity represents the normalized median intensity for each well position. **(c)** A scatter plot displaying the averaged median intensity values over three replicates ranked in ascending order, with normalized median intensity for each strain and corresponding standard deviation (StdDev) as error bars.

### Utilization of a Genetically Encoded Affinity Assay for Visualizing Protein Localization

Tag size raises a concern regarding the interference of such an appendage with protein structure and function. However, smaller tags usually do not enable *in vivo* visualization. Therefore, we developed a system for monitoring the localization of the N’ HA-tagged proteins in live cells. To this end, we mated the HA library with a strain expressing a single-chain variable fragment that specifically binds the HA tag (scFv_HA_) fused to a fluorescent protein (Tsirkas et al., 2022; Zhao et al., 2019) . The scFv_HA_ was fused to a yeast-codon optimized yomScarlet-I3 fluorescent protein, which is a bright and rapidly maturing protein (Gadella et al., 2023). The scFv was expressed under the control of the inducible Z3 promoter on the background of the Z3 transcription factor (Z3TF) that controls the Z3 promoter (McIsaac et al., 2013; Ohira et al., 2017) in a BY4742 (Mat α) strain (Figure 4a). Upon addition of β-estradiol to the media, the Z3TF binds to the Z3 promoter, and induces the expression of scFv_HA_-yomScarlet-I3 (from here on scFv_HA_-Scarlet), thereby enabling the visualization of HA-tagged proteins. Having an inducible expression system allows the N’ HA proteins to fold, assemble and function whilst only having the small tag, but be visualized, on demand, as the scFv is expressed.

**Figure 4.**
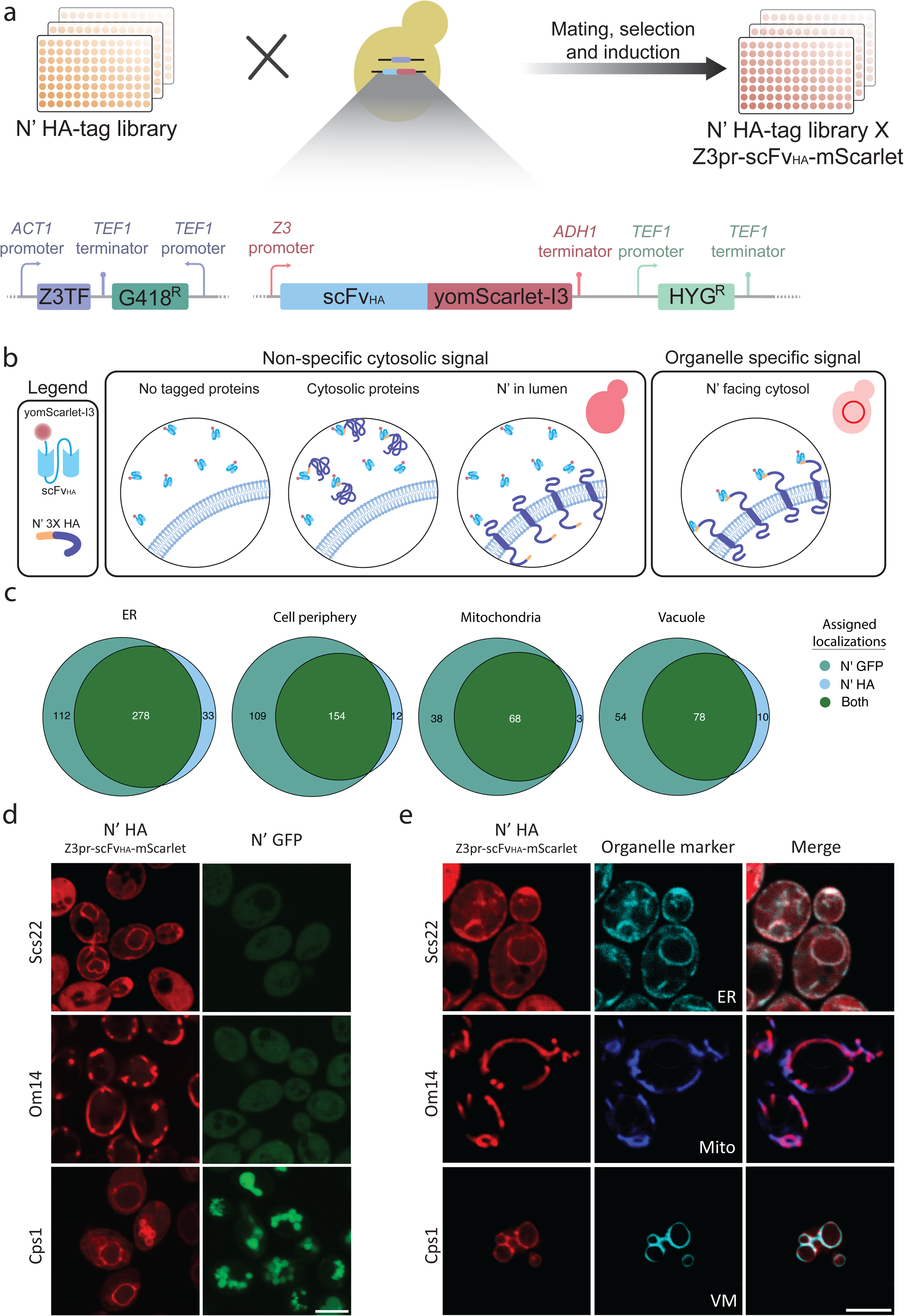
Visualization of protein localization using a genetically encoded affinity assay. (a) Schematic representation of the creation of the diploid library: The HA-tagged protein library was crossed against a strain expressing scFv_HA_-yomScarlet-I3 (mScarlet) under the inducible Z3 Transcription Factor (TF). Upon β-estradiol induction, the scFv_HA_-mScarlet was expressed in the cytosol. Diploid strains were imaged by confocal microscopy to detect localized yomScarlet-I3 fluorescent signal. **(b)** Illustration of expected phenotypes for different N’ localization of HA-tagged proteins using the scFv_HA_-mScarlet, Z3TF system. Untagged proteins, cytosolic proteins, and proteins with N’ facing lumens of organelle should show a cytosolic signal as the scFv_HA_-mScarlet is dispersed throughout the cytosol. Membrane proteins with cytosolic N’-HA will interact with the scFv_HA_- mScarlet and therefore present as a localized fluorescent signal of an organelle-specific nature. **(c)** Venn diagram of proteins exhibiting organelle specific signal in the *NOP1-*GFP and the HA x sc-Fv_HA_ libraries were analyzed for discrepancies in their localization. 65.7% of the GFP-tagged proteins that are localized to the Endoplasmic Reticulum (ER) also show a similar signal when tagged with HA. Whilst 56% correlate between GFP and HA for the cell periphery, 62.4% for mitochondria and 54.9% for the vacuole/vacuole membrane. **(d)** Confocal microscopy images of representative strains that exhibited clear localizations to various organelles in the HA x sc-FV_HA_ library (red) despite not being clearly localized when tagged with GFP (green): Scs22 has a clear ER localization, Om14 a mitochondrial one, and Cps1 vacuolar membrane. Diploid strains of both backgrounds were induced with β-estradiol and imaged after 3 hours alongside their counterparts in the *NOP1-GFP* library. Scale bar = 5µm. **(e)** Confocal microscopy images of the representative strains expressing organellar markers (cyan) show full colocalization of the HA x sc-FV_HA_ strains (red) with the respective fluorescently labeled organelles. Sec63-mNeonGreen was used as a marker for the ER, MitoView405 was used as a mitochondrial marker and Vph1-mNeonGreen was used as a vacuole membrane (VM) marker. Scale bar = 5µm.

Since the scFV_HA_-Scarlet is expressed in the cytosol it is not possible to use it to visualize cytosolic proteins nor proteins sequestered inside organelle lumens. However, membranal proteins whose N-terminus is facing the cytosol will bind the fluorophore to the surface of their respective organelles of residence, resulting in a localized fluorescent signal specific to the resident organelle (Figure 4b).

To calibrate the optimal conditions for induction of the scFv_HA_-Scarlet, we measured its expression in the presence of different β-estradiol concentrations (0, 50, 100, and 1000 nM) at different times and chose 100nM at a time range of 3-4 hours for further analysis to allow Scarlet signal detection but prevent saturating (Supplementary figure S2a and b). To validate these conditions, we manually picked from the library strains with N’-HA proteins from different cellular compartments with known topologies of N-termini facing the cytosol [Alg7 (endoplasmic reticulum (ER)), Gem1 (mitochondria), Pst2 (cell periphery), Tpd3 (cytosol), Zrc1 (vacuole)], and mated them with the scFv_HA_-Scarlet strain. We then imaged these strains at 150 and 180 minutes after addition of 100nM beta-estradiol, validating their visualization at the expected organelle (Supplementary figure S2c).

Using these optimized conditions, we then mated the entire HA library with the scFv_HA_-Scarlet strain and screened the diploid library for labeled proteins that exhibited organelle-specific signals. These proteins were compared to the *NOP1*-GFP library to identify any proteins whose localization may be better visualized by our system (Figure 4c, Supplementary table S2). We focused on strains that displayed organelle-specific localization in our HA library and removed proteins with signal peptides (SP) and mitochondrial targeting sequences (MTS) since they would reside in the lumen/matrix of organelles clearly not compatible with our system. Our findings indicate that 65.7% of GFP-tagged proteins in the ER showed a similar signal when tagged with HA. Additionally, 56% of proteins in the cell periphery, 62.4% of mitochondrial proteins and 54.9% of vacuolar ones had similar localization when tagged with the HA tag.

Interestingly, 3-8% of the localizations observed in the HA library were novel, suggesting unique advantages of the HA library under specific conditions compared to the GFP library. This observation was manually validated by comparing strains showing new localizations in the HA library with those in the *NOP1*-GFP library. For example, Scs22, Om14, and Cps1 exhibited unique organelle-specific localization in the HA library. Scs22 is a well characterized contact site protein, homologous to the mammalian VAP proteins, which is anchored to the ER membrane via a C’ transmembrane tail anchor (Manford et al., 2012). Om14 is a validated mitochondrial outer membrane protein (Burri et al., 2006) that could not be visualized using an N’ GFP tag but is now clearly visualized in mitochondria with the HA epitope (Figure 4d and e). Cps1 is a vacuolar carboxy peptidase with an N’ transmembrane domain that is first targeted to the ER membrane and then transported to the vacuolar membrane. The C’ of Cps1 faces the lumen, where it is cleaved, releasing the catalytic subunit of the protein from the membrane to the vacuole lumen (Spormann et al., 1992). In the HA library, we found that the Scarlet signal of Cps1 is clearly marking the vacuolar membrane, while in the *NOP1*-GFP strain, the GFP signal localizes to the vacuolar lumen, indicating that the smaller HA tag allows for a more accurate depiction of Cps1’s localization (Figure 4d and e). These results underscore the benefits of using a smaller tag, such as HA, for precise experimental applications.

Building on the success of the scFv_HA_ system we developed an expansion of the scFv anti-HA toolkit that incorporates organellar markers, photoconvertible (mEOS3.1), and photo-switchable (ffDronpa) proteins (Supplementary figures S3a,b, and c). These advanced fluorescent proteins, which change their fluorescence upon light exposure, allow precise temporal and spatial tracking of protein dynamics significantly improving the versatility and depth of analysis possible with the library toolset.

## Discussion

In this study, we developed an N’-HA yeast proteome-wide library, addressing the limitations associated with larger tags utilized in existing libraries. The small HA tag offers significant advantages by reducing the risk of interfering with protein folding, stability, assembly, localization or activity, while still enabling effective detection and functional assays. The utilization of the SWAT strategy allowed the creation of the N’ HA-tagged library, with high integration efficiency and proteome-wide coverage.

To demonstrate the versatility of the HA library, we showed its utility in the analysis of protein molecular weight and post translational modifications rapidly and systematically. Our ability to quantify protein abundance enables the discovery of factors that control protein expression beyond transcriptional regulation. Finally, the visualization of the N-HA protein by the scFv approach enables the *in vivo* characterization of the target proteins at its designated organelle. The use of the small HA tag, while less disruptive than larger tags, can still presents challenges in specific contexts. While the N’ HA-tagged protein bound to scFv_HA_-Scarlet system was effective for visualizing proteins in organelle membranes with cytosolic N-termini, it is not suitable for detecting proteins that are completely cytosolic or with the opposite topology where the N-terminus is inaccessible to the scFv_HA_. Two exceptions for this are proteins targeted to the peroxisome or the nucleus lumen, since their membrane import mechanisms can accommodate larger complexes (Meinecke et al., 2016; Rout and Aitchison, 2001). Indeed, by utilizing the library, we visualized luminal peroxisomal proteins, including Pex8, Pxp1, and Pcs60 and luminal nuclear proteins, such as Net1, Rsc6, and Nop10 in the correct cellular compartments.

Our reliance on a single, constitutive promoter in the HA library does not account for natural promoter-specific expression variability, which could limit its application in studying dynamic gene regulation across different conditions. However, since each library has its benefits and limitations, we foresee that this library will support a niche of assays that will benefit from this design. For other uses it is possible to use the SWAT approach to efficiently create a “Seamless” library where the HA tag is fused at the N’ but under the natural endogenous promoter (REF: Yofe et al, 2016) or a C’ tagged library (Ref: Meurer et al).

While the HA library can be used to study any protein of choice, it is probably most powerful for studying very small ones where the large FP tag is a dramatic appendage. Recent insights into the role of small open reading frames (sORFs) provides additional dimensions to consider in proteomics research. sORFs, encoding for proteins under 100 amino acids long, are often overlooked due to their size. However, they encode for small peptides that can have critical biological functions, including roles in cellular signaling, stress responses, and translation regulation (Couso and Patraquim, 2017; Kastenmayer et al., 2006). Due to their small size, it is critical minimize the size of appendages fused to them. Therefore, the N’-HA library provides an opportunity to systematically study these small peptides, potentially uncovering new functions or regulatory mechanisms. Upon analysis, 319 out of the 365 sORFs from the original SWAT library were retained in our final library, in line with the measured survival rate (supplementary Table S1). Overall, our findings highlight the advantages of a proteome-wide N’ HA-tagged yeast library as a valuable tool for functional genomics. By minimizing the impact on protein structure and localization, the HA tag provides a more accurate representation of native protein behavior, enabling systematic analyses that were previously challenging with larger tags. The applications demonstrated in this study, such as glycosylation detection, quantification of protein abundance, and *in vivo* visualization of membrane proteins illustrate the breadth of possibilities offered by this comprehensive library. Moving forward, the N’-HA library holds promise for further discoveries in yeast biology and may serve as a model for similar libraries in other organisms.

## Materials and methods

### Yeast strains and plasmids

All yeast strains used in this study are listed in supplementary table S3. Strains were constructed using the lithium acetate-based transformation protocol (Gietz and Woods, 2002). All plasmids used are listed in supplementary table S4 and primers listed in supplementary table S5.

### Yeast library generation

SWAT library generation was performed as described (Weill et al., 2018). Briefly, a RoToR array pinning robot (Singer Instruments) was used to mate the parental NP tag GFP SWAT library with the required donor strain (Supplementary table S3) and carry out the subsequent sporulation, and selection protocol to generate a haploid library selected for all the desired features. Growth of the library on YPGalactose (2% peptone, 1% yeast extract, 2% galactose) was used to induce SceI-mediated tag swapping, and subsequent growth on SD containing 5-fluoroorotic acid (5-FOA, Formedium) at 1 g/l, and required metabolic and antibiotic selections were used to select for strains, which had successfully undergone the SWAT process.

### Diploid strain generation

The N’ HA library was grown on SD supplemented with NAT at 30°C overnight and the scFv_Anti HA_ -yomScarlet-I3, Z3TF donor strain (Supplementary table S3) was grown on YPD supplemented with geneticin (G418) and Hygromycin B (HYG). Both were replicated onto a YPD plate and grown overnight at room temperature (RT). The mated strains were then replicated onto SD with monosodium glutamate (MSG) plates supplemented with NAT, G418, and HYG and grown overnight at 30°C. This step was repeated once more to select diploid strains containing the combination of desired traits.

### Protein extraction and SDS-PAGE for library validation

5ml of cells at 0.5OD_600_ were collected by centrifugation at 3,000P*g* for 3Pmin, washed with 1Pml of double-distilled water (DDW), resuspended in 200Pμl lysis buffer containing 8PM urea, 50PmM Tris pH 7.5, and complete Protease Inhibitors (Merck), and lysed by high-speed bead beater with glass beads (Scientific Industries) at 4°C for 10Pmin. 25Pμl of 20% Sodium Dodecyl Sulfate (SDS) was added before incubation at 45°C for 15Pmin. The bottom of the microcentrifuge tubes was then pierced, loaded into 5Pml tubes, and centrifuged at 4,000P*g* for 10Pmin to separate the lysate from the glass beads. The flow-through collected in the 5Pml tubes was transferred to a fresh 1.5Pml microcentrifuge tube and centrifuged at 20,000P*g* for 5Pmin. The supernatant was collected, and 4× SDS-free sample buffer (0.25PM Tris pH 6.8, 15% glycerol, and 16% Orange G containing 100PmM DTT) was added to the lysates, and incubated at 45°C for 15Pmin. Protein samples were then separated by SDS–PAGE using a 12% polyacrylamide gel and then transferred onto a 0.45Pμm nitrocellulose membrane (Pall Corporation) using a Trans-Blot Turbo transfer system (Bio-Rad). Membranes were blocked in bovine serum albumin Buffer (BSA) in phosphate-buffered saline (PBS) solution for 30Pmin at RT, incubated over-night at 4^0^c with rabbit anti-Histone H3 (ab1791, 1:5,000; Abcam) and mouse anti-HA (#901502 1:1000, BioLegend) diluted in a 2% wt/vol BSA/PBS solution containing 0.01% NaN3. After washing 3 times in TBST buffer, membranes were then probed with secondary goat anti-rabbit-IRDye680RD antibody (#ab216777; Abcam) and 800CW Goat anti-mouse IgG (#ab216772; Abcam), both diluted 1:10,000 in 3% wt/vol milk/TBST solution for 1Ph, at RT. Blots were washed and imaged on the LI-COR Odyssey Infrared Scanner (ODY-2064).

### Library coverage analysis

Images of agar plates of the N’ HA library were taken by a Canon digital camera. The images were subsequently processed by SGAtools v2.3 (Wagih et al., 2013), to measure the size and shape of each colony and established a minimum size threshold for inclusion as a valid colony.

### Post-Translational Modification Analysis

Yeast strains were grown overnight in YPD media containing NAT at 30°C. The following day, the strains were back diluted to an OD_600_= 0.2 and grown at 30°C for 4 h. Next, 2.5 OD_600_ of yeast cells were harvested, washed once in DDW, resuspended in 0.1 M NaOH, and incubated for 5 minutes at RT. The cells were then centrifuged at 3000 *g* for 3 minutes. The pellets were subsequently assayed for glycosylation by PNGase-F kit (New England Biolabs, P0704S) according to the manufacturer’s protocol and resolved by SDS-PAGE.

## High Throughputs Quantitative dot-blot

### Protein extraction

Yeast strain colonies were inoculated into a 1 ml polypropylene 96 deep-well plate, filled with 400 µl of YPD medium supplemented with 200 µg/ml of NAT as a selective agent. After inoculation, the plate was sealed with a breathable cover (AeraSeal BS-25, Excel scientific) and placed in a shaker incubator set to 30°C with shaking at 650 rpm for overnight growth. The following day the 96-well plate was spun down using a swing-out rotor (Eppendorf model 5810R) at 3000 *g* for 5 minutes. The resulting pellets (roughly two OD per sample) were washed once with TE buffer and stored at -20 °C until further use. Subsequently, samples were resuspended in 220 µl of Protein Extraction Buffer (8M Urea, Tris-HCl pH 7.5). To break open the yeast cell wall, another 1 ml, deep 96-well plate was customized by drilling a 1 mm diameter hole at each of its well’s bottom. Then, the punctured wells were carefully sealed with an aluminum sticker (PCR-AS-200, Axigen) before being filled with ∼100 µl of acid-washed glass beads (Merck cat. G8772) and 100 µl of Urea-suspended samples. Next, the wells were sealed with a second aluminum sticker, and the plate was mounted on a vortex with a flat adapter (Scientific Industries, Part No.504-0235-00) and vortexed for 15 minutes at maximum vibration frequency (3220 rpm) in a cold room. Then the bottom aluminum sticker was peeled off just before securing it atop a new 96-well, fully skirted PCR plate with adhesive tape. The tandem plates were spun at 800 *g* for 3 minutes using a swing-out centrifuge to transfer the extracted sample into the lower PCR plate. A new PCR plate was set up for the final preparation by adding 15 µl of 4x Laemmli buffer (supplemented with DTT) to each well. Then, 45 µl of the extracted samples were transferred to this plate, mixed by pipetting with the Laemmli buffer, sealed with a PCR sticker, and heated to 95 °C for 10 minutes.

### Dot-Blotting

For dot blotting, a nitrocellulose membrane, soaked in protein transfer buffer (Bio Prep, TB192), was placed on a sheet of Whatman blotting paper (No.3) laid atop the bottom part of the Dot-blotter manifold. After soaking, the manifold’s upper part was assembled and secured with clips. Then 35 µl of each sample was loaded into its designated position in the manifold using a multichannel pipette while the manifold was connected to a vacuum suction. The dotted membrane was then subjected to a Trans-Blot Turbo transfer system (Bio-Rad) to immobilize the protein samples, with a transfer program set to run for 7 minutes at 2.5A. Following immobilization, primary (anti-HA, Roche, 11867423001) and secondary (IRDye 800CW Goat anti-rat, Li-Cor, 92632219) antibodies were applied according to standard Western blotting protocols.

### Image acquisition

The near-infrared fluorescence signals of the HA-tagged proteins were acquired from the dot-blot membrane using an Odyssey infrared imaging system (ODY-2064, Li-Cor Biosciences). The signal intensity was normalized by BCA-protein quantification assay (Thermo scientific, 23225). Image captured through Odyssey software (v3.0.21) with standardized scanning parameters, including a 169 µm resolution and intensity settings 5.0 for the 800 nm channel. The images were then exported in TIFF format to preserve the original pixel intensity values for subsequent analysis.

### Signal enhancement

Prior to quantification, we systematically followed a quality control workflow to ensure robust and reproducible measurements. Dot-blot images were individually inspected for potential technical artifacts or poor signal quality that might compromise accurate quantification. Notwithstanding our high-quality imaging, we still implemented a two-step background correction in ImageJ/Fiji (v1.54f) to account for slightly uneven illumination: (a) a noise reduction using a Gaussian blur filter (σ = 2 pixels) and (b) a local background estimation of the blurred image using the rolling-ball algorithm (radius = 35 pixels) which was subsequently subtracted from the original image.

### Signal quantification

Our custom Python software (ht-qdotblot) enables semi-automated quantification of signal intensity across the 96-well format. The software implements a grid-based quantification approach, where users initially register three reference points corresponding to the centers of the plate corners (wells A1, A12, and H1). The software then automatically overlays a virtual grid that can be fine-tuned through adjustments of well radius, spacing, and positioning. A grid-based approach allows for the selection of consistent region-of-interest across multiple images and experimental conditions. For each well, the software returns multiple intensity-based measurements, including median, mean, standard deviation, mode, and minimum/maximum value. The data can be exported in CSV format for further analysis and visualization. Our GitHub repository contains the software source code and provides a guided tutorial for installation and usage on an example image (https://github.com/benjamin-elusers/HT-qDotblot).

### Sample normalization

To account for technical variations in protein loading and transfer efficiency, we employed a normalization strategy. Total protein content was quantified using the BCA assay (thermoscientific, Cat#23225), providing a normalization factor related to actual protein concentration in each well. Furthermore, we subtracted from the normalized median intensity signals, the background values of two internal controls (buffer-only and untagged yeast). The final sample integrated intensities were calculated as the mean of background-subtracted normalized median intensities across replicates, which were scattered on the plate to avoid any spatial bias in the quantification. For the 32 samples, we report the individual integrated intensities along with their mean ± SEM from three independent replicates.

### Confocal microscopy

For high-throughput screening of the full library, cells were moved from agar plates into liquid 384-well plates using the RoToR bench-top colony arrayer (Singer Instruments). Liquid cultures were grown overnight in synthetic medium with 2% glucose (SD) in a shaking incubator (LiCONiC Instruments) at 30°C. A Tecan freedom EVO liquid handler (Tecan), which is connected to the incubator, was used to back-dilute the strains to _∼_P0.25POD_600_ in plates containing SD with 100nM β-estradiol. Plates were then transferred back to the incubator and were allowed to grow for 4Ph at 30°C to reach logarithmic growth phase. The liquid handler was then used to transfer strains into glass-bottom 384-well microscope plates (Azenta Life Sciences) coated with 0.25 mg/ml Concanavalin A (Sigma-Aldrich) to allow cell adhesion. Wells were washed twice in SD to remove floating cells and reach a cell monolayer. Plates were then automatically moved by a KX-2 robotic arm (Peak Robotics) into an automated inverted spinning disk microscope system (Olympus) and imaged using a 60x air lens (UPlanFLN, NA 0.9).

For all other microscopy-based figures, images were obtained using an automated inverted fluorescence microscope system (Olympus) containing a spinning disk high-resolution module (Yokogawa CSU-W1 SoRa confocal scanner with double micro lenses and 50 μm pinholes).

Several planes were recorded using a 60x oil lens (NA 1.42) and with a Hamamatsu ORCA-Flash 4.0 camera. Fluorophores were excited by a laser and images were recorded in three channels: GFP (excitation wavelength 488 nm, emission filter 525/50 nm), mCherry / mScarlet / FM™4-64 (excitation wavelength 561 nm, emission filter 617/73 nm) and MitoView™405 (excitation wavelength 405 nm, emission filter 447/60). Image acquisition was performed using scanR Olympus soft imaging solutions version 3.2. Images were transferred to ImageJ, for slight contrast and brightness adjustments to each individual panel. Images were manually inspected using Fiji-ImageJ software (Schindelin et al., 2012).

## Acknowledgments

We thank Rosario Valenti and Olga Beresh for critical reading of the manuscript and helpful suggestions. We thank prof. Michael Toledano for the idea to create this library. We thank Dr. Yoav Peleg for help with plasmid generation.

This work was supported by a Chan Zuckerberg Initiative (CZI) grant (2023-331952) and by the European Union ERC CoG (OnTarget 864068) as well as by the Institute for Environmental Sustainability (IES) at the Weizmann Institute of Science. The robotic system in the Schuldiner lab was purchased through the kind support of the Blythe Brenden-Mann Foundation. MS is Incumbent of the Dr. Gilbert Omenn and Martha Darling Professorial Chair in Molecular Genetics.

## Author Contributions

Conceptualization: DB, OK, IT, AA, MS, ES, BD Data curation: DB Formal analysis: DB, OK, YA Funding acquisition: MS, AA Investigation: DB, OK Methodology: ES, BD, IT Software: BD Supervision: OK, AA, MS Visualization: DB Writing – original draft: DB, OK, MS Writing – review & editing: All authors

## Supplementary figure legends

**Supplementary Figure 1:** Verification of the N’ HA-tagged yeast library using Western blot analysis. Western blots of selected HA-tagged yeast strains to verify successful, in frame, integration of the HA tag. Each lane represents a different HA-tagged protein from the library, with anti-HA antibodies used to detect the presence and correct size of the tagged proteins. Control lanes include a non-HA-tagged yeast strain as a negative control (BY4741) to confirm specificity of the anti-HA antibody. Molecular weight (MW) markers indicate size ranges for each HA-tagged protein. Bands at expected sizes confirm that most proteins are correctly tagged and expressed. Anti histone H3 was used as a loading control. Protein names in red indicate lack of a clear band.

**Supplementary Figure 2:** Calibration and optimization of the scFv anti-HA system for imaging. (a) Plot of intensity (arbitrary units, AU) over time (minutes) in yeast strains containing the scFv_HA_-Scarlet system induced with β-estradiol at concentrations of 50nM, 100nM, and 1000nM. A strain without β-estradiol was used as a control (none). The plot highlights the optimal time and β-estradiol concentration (indicated with brackets and a camera icon) for achieving a coherent visualized signal (100nM, ∼180-250min). **(b)** Representative confocal images of yeast strains with the scFv_HA_-mScarlet system at various time points and β-estradiol concentrations from (a). Images demonstrate increasing fluorescence intensity with higher concentrations and longer induction times. Scale bar = 5 µm. **(c)** Representative confocal images of diploid yeast strains (N’-HA + scFv_HA_-mScarlet) with HA-tagged proteins localized to various organelle membranes: Alg7 to the endoplasmic reticulum (ER), Gem1 to mitochondria, Pst2 to the cell periphery, Tpd3 in the cytosol, and Zrc1 to the vacuole membrane. Images were taken at 0, 150, and 180 minutes after β-estradiol induction to monitor changes in signal intensity. The fluorescent signals confirm the accurate localization of HA-tagged proteins across these compartments. Scale bar = 5 µm.

**Supplementary Figure 3:** Expansion of the scFv anti-HA toolkit with organellar markers, photoconvertible and photo-switchable proteins. **(a)** Confocal images of diploid yeast strains containing the scFv_HA_-mScarlet system (red) alongside genome-integrated organelle markers (green), as used in figure 4e: Vph1 (vacuole membrane, VM) and Sec63 (endoplasmic reticulum, ER). Scale bar = 5 µm. **(b)** Confocal images of a yeast strain containing scFv_HA_-mEOS3.1, a photoconvertible fluorophore. The images show the photoconversion from green to red after illumination by a 405nm laser, expanding the toolkit’s versatility for dynamic protein tracking. Scale bar = 5 µm. **(c)** Confocal images of a yeast strain containing scFv_HA_-ffDronpa, a photo-switchable fluorophore, imaged at 488 nm in both “on” and “off” states. The “on” state was reactivated following excitation with illumination at 405 nm wavelength. These images demonstrate that scFv_HA_-ffDronpa can be efficiently toggled between fluorescent states, allowing controlled imaging in live cells. Scale bar = 5 µm.

## Notes

### Competing Interest Statement

The authors have declared no competing interest.

