## Supplementary figures and images for "Creation and Validation of a Proteome-Wide Yeast Library for Protein Detection and Analysis"

### Supplemental figures

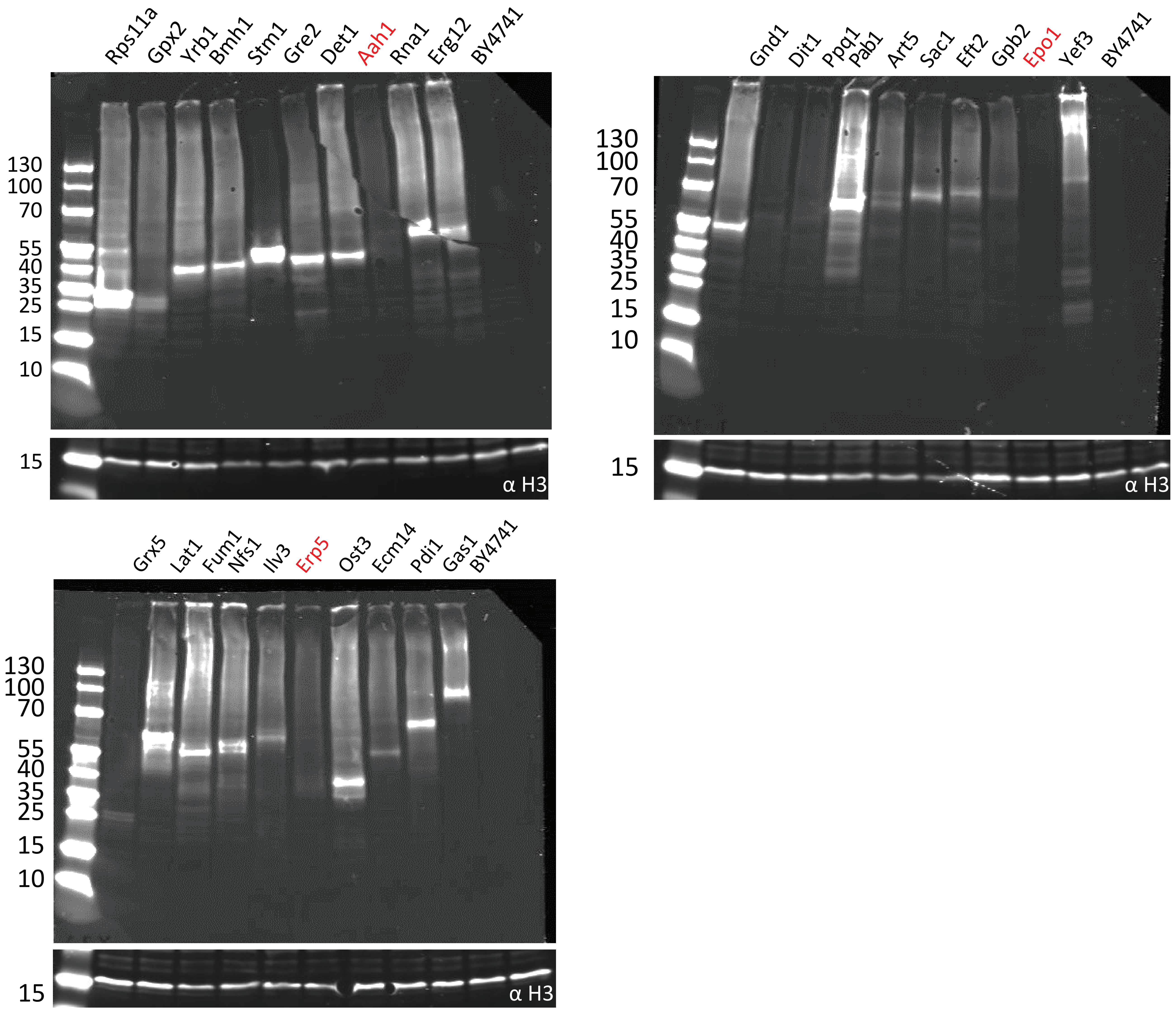

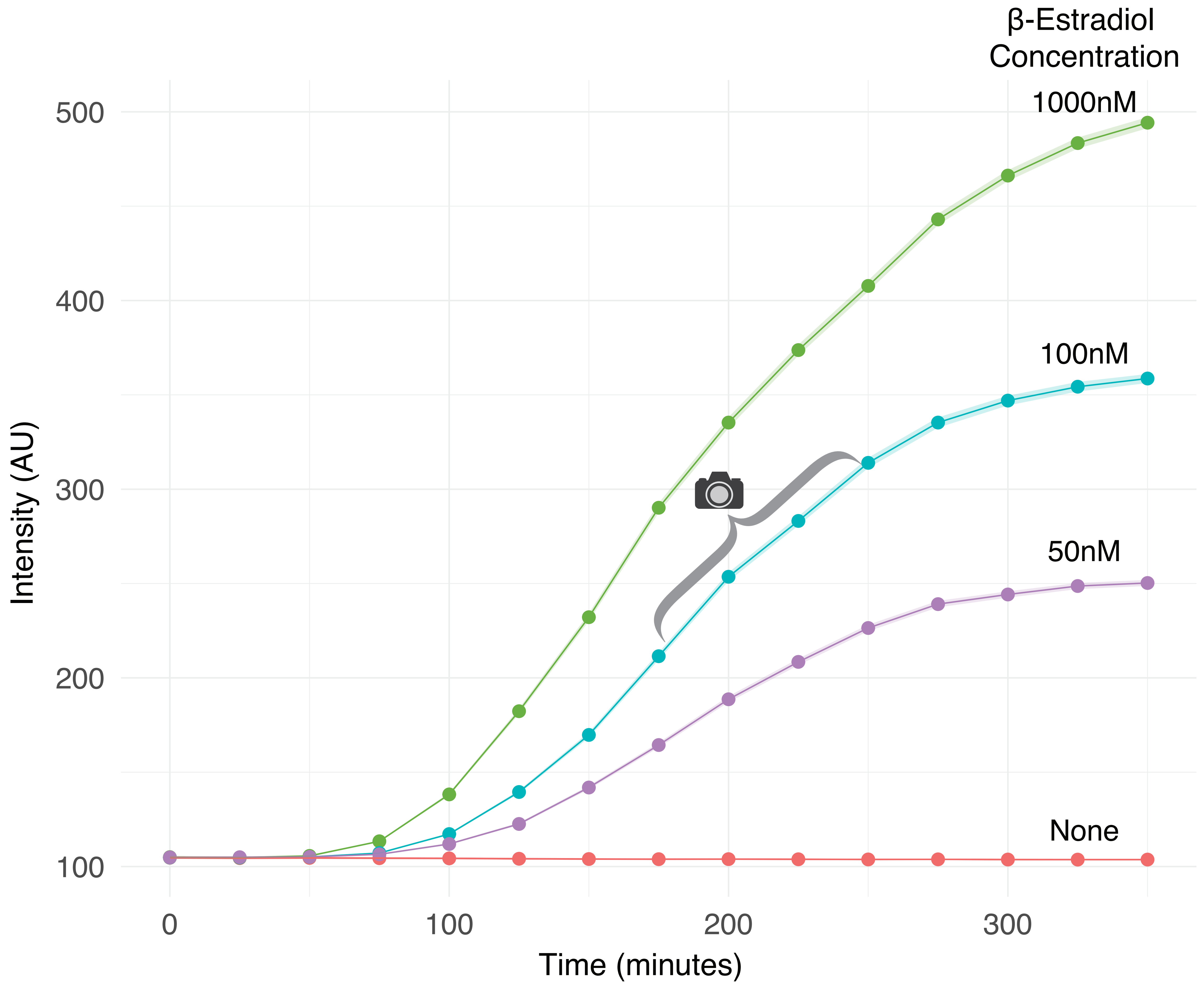

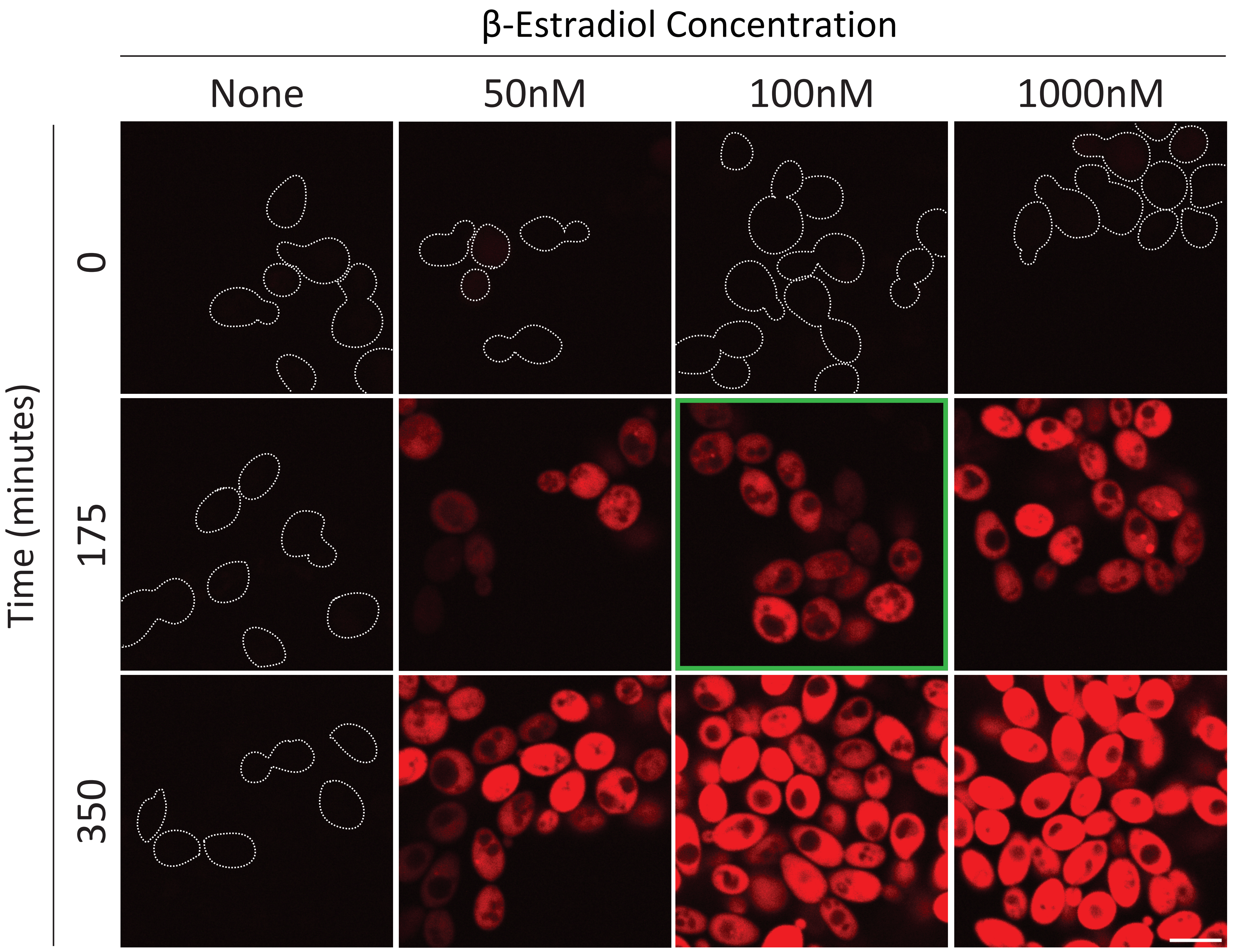

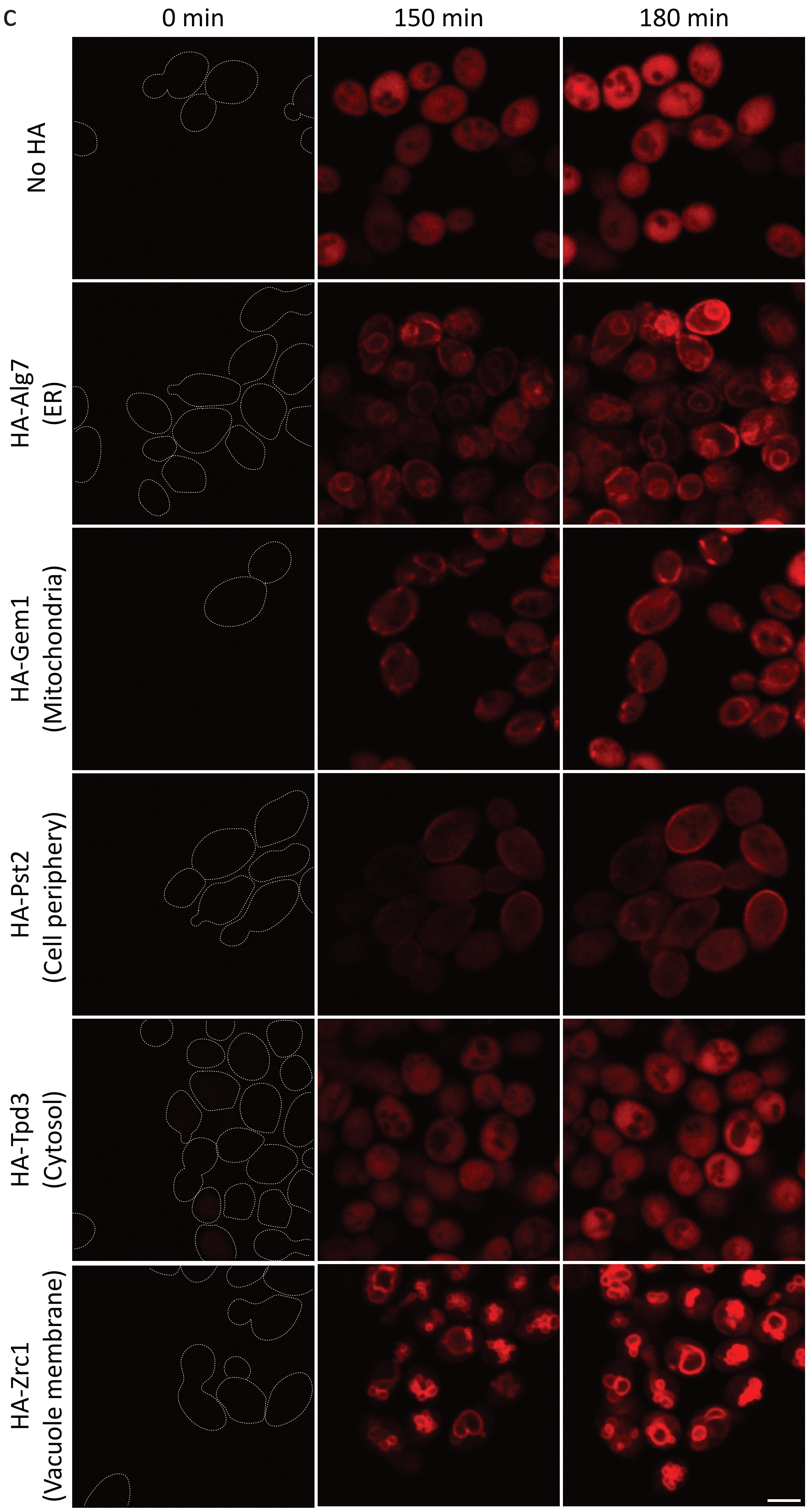

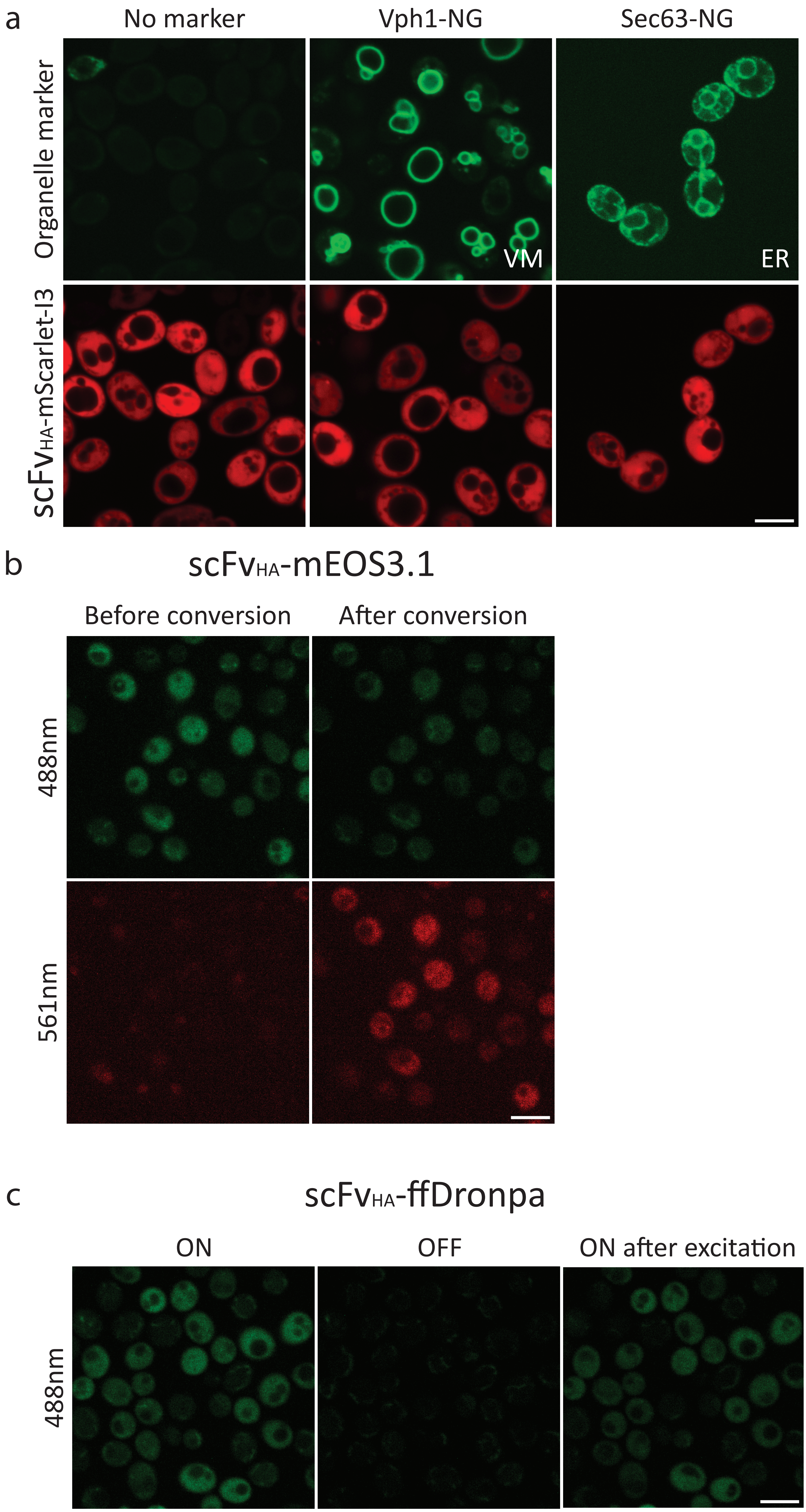
